# A compact prism-based microscope for high sensitive measurements in fluid biopsy

**DOI:** 10.1101/2024.08.13.607708

**Authors:** Laura Perego, Caterina Dallari, Chiara Falciani, Alessandro Pini, Lucia Gardini, Caterina Credi, Francesco Saverio Pavone

**Author notes:** Corresponding Author Caterina Credi. These authors contributed equally (Laura Perego, Caterina Dallari).

## Abstract

The increasing demand for sensitive, portable, and cost-effective disease detection methods has raised significant interest in the development of biosensors for rapid and early-stage diagnosis, population mass-screening, and bedside monitoring. Although a certain number of sensing devices with high sensitivity has experienced considerable progress, their practical application has been hindered by challenges in replicating instrumental systems, obtaining rapid and multiplexed signals, and the high cost of manufacturing. In response, we present a streamlined prism-based Total Internal Reflection system which, in combination with surface functionalization techniques on gold nanoparticles, is capable of facilitating Evanescent Wave scattering for the highly sensitive and rapid detection of specific analytes in synthetic liquids and real human samples. The system achieves a remarkable limit of detection of 1 fg/mL concentration for the targeted pathological biomarker. Our innovative design addresses the limitations of existing technologies by reducing costs, minimising overall size, and ensuring swift biofluid analysis with remarkable sensitivity. To validate its efficacy, we conducted scattering experiments in synthetic and human serum samples, exploiting functionalized AuNPs to recognize bacterial lipopolysaccharides as biomarkers for sepsis disease. The cohesive integration of these techniques hopefully makes this biosensing setup a promising candidate for potential clinical deployment, meeting the pressing requirements for rapid personalised diagnosis, large-scale population screening and bed-monitoring.

## INTRODUCTION

In recent years, liquid biopsy has emerged in the clinical scenario as a powerful and straightforward non-invasive diagnostic alternative to the “gold-standard” assays and to tissue biopsy, which represents the current methods for disease detection through histological analysis or genetic evaluation ^1^. Indeed, standard biopsy methods present some disadvantages being highly invasive, risky, and impractical, especially for deep-seated lesions, and they may require long and expensive analytical procedures and equipment. The emergence of liquid biopsy as an alternative to standard biopsy methods raises the need for novel technologies. Thus, a big effort has been directed towards the development of new highly-sensitive technologies enabling the analysis of body fluids and the detection of abnormal concentration/alteration of pathological biomarkers with improved sensitivity and specificity ^2^. Recurrent monitoring of biomarkers could give real-time knowledge of the status of the patient and help both the prevention screening, before the disease progression, and the treatment decisions by gathering information on the dynamics of the pathology ^3,4^. In this scenario, total internal reflection (TIR) spectroscopy has been exploited as a non-destructive, selective and extremely sensitive method to analyse adsorption phenomena and biomolecular interactions ^5^. In particular, TIR microscopy exploits such phenomenon occurring at the interface between two materials with different refractive index: when a collimated beam impacts the interface between the glass coverslip and the water of the sample solution with an angle of incidence higher that the critical angle, total internal reflection occurs, following Snell’s law, and the incident beam is totally back reflected from the glass surface through the incidence optical path (Figure 1, inset). This phenomenon is accompanied by the propagation of an Evanescent Wave (EW), characterised by the same wavelength of the incident beam and with an intensity that decays exponentially within a characteristic distance of about 100-200 nanometres inside the second medium (i.e. the biofluid to be tested) (Figure 1). The low penetration depth of the EW can be exploited to limit the excitation volume to a very small portion of the sample, thus reducing the background signal coming from deepest regions of the sample, while retaining a wide field configuration^6^. In this configuration, only the molecules located in the EW region are excited and the contrast is enhanced, thus improving the detection sensitivity down to the single molecule level ^7^. Depending on the type of labels or probes linked to the analytes (fluorophores, metal particles) different optical signals can be detected, spanning from fluorescence to scattering (both elastic and inelastic) ^7–9^. The choice of TIR scattering microscopy, using metallic nanoparticles as probes, is encouraged by the characteristic NPs large scattering cross-section. Self-assembly of gold nanoparticles (AuNPs) on the analytes immobilised in the EW region can be exploited to further enhance the scattered optical signal up to 6-8 orders of magnitude, enabling for nanometric precision characterization of single particles ^10,11^. Moreover, in order to improve the specificity of these detection systems towards specific analytes dispersed in body fluids, it is possible to engineer the chemistry of the surface of the NPs and to induce specific NPs-capturing of the biomarker of interest, thus enabling their selective recognition ^12^. This strategy allows to directly detect biomarkers in unprocessed biological matrices, thus avoiding pre-processing procedures or separation steps that could alter the chemistry of the samples to be analysed and extend the time for analysis. Furthermore, thanks to the recent technological advances in micro- and nanofabrication processes, EW biosensors could also strongly benefit from integration with lab-on-chip (LoC) microfluidic devices, thus enabling for smart handling, efficient mixing, sorting or separation of few microliters of samples, as well as for easier implementation of multiplexing analysis ^13,14^.

**Figure 1:**
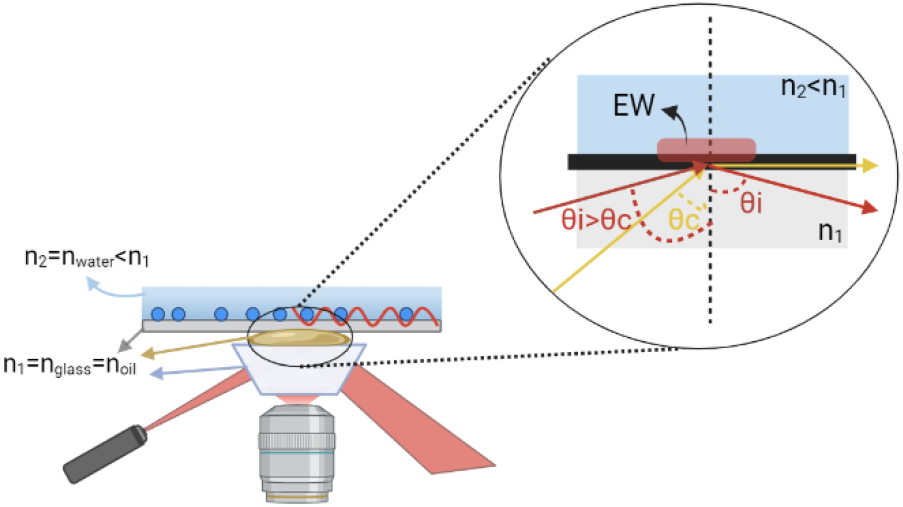
Graphical scheme depicting the optical coupling between the glass microprism and the glass coverslip through the usage of immersion oil, which presents the same refraction index of glass. A refraction index mismatch occurs at the interface between the coverslip and the sample buffer. As a consequence, TIR occurs at this interface, generating a EW which propagates within the sample buffer. In the inset, the behaviour of light rays at the interface between two media with different refraction indexes, according to Snell’s law, is shown.

Here we report on the full development of a modular low-cost bench-top TIR detection system characterised by high portability to be coupled with an optical biosensor for the screening of analytes in biofluids, and on their further miniaturisation within LOCs. The combination between microfluidic chambers containing NPs engineered for the selective recognition of biomarkers of interest and the TIR scattering setup forms an advanced optical biosensor platform which is very versatile, allowing for the incorporation of an almost limitless range of biorecognition probes precisely and robustly conjugated to the sensor by covalent surface chemistry approaches. As a proof of concept, following previous works published by our group ^15^, the surface of NPs has been fully functionalized with ad-hoc synthesised nonnatural antimicrobial peptides, engineered to specifically recognize bacterial lipopolysaccharides (LPS). Changes in blood-LPS levels are signs of severe infections and sepsis ^16–18^. Thus, early and precise diagnosis with liquid biopsy can be crucial in improving patients’ prognosis ^19,20^. First, the peptide/NP ratio was optimised to achieve a complete covering of the gold surface, yet ensuring the retaining of the colloidal stability at the same time ^21–23^. Then, a low-cost bench-top TIR microscope was successfully designed and realised, which exploits EW scattering to reveal analytes in fluids with sensitivity and specificity comparable to standard bulk TIR microscopes, but with a strong reduction of costs and dimensions. The proposed synergic combination of advanced EW-based compact setup and ad-hoc functionalized NPs for the screening of LPS in real serum samples will pave the way for possible applications in clinics for early-stage diagnosis, with an advanced level of automatization and high-throughput.

## 2. METHODS

### 2.1. Implementation of the portable optical detection system

Since our final goal was the collection of the scattering signal coming from gold nanoparticles specifically coupled with the analytes to be detected, and since these measurements were intended for early-stage detection of diseases, a straightforward and affordable optical reader for the scattering signal was required. To achieve this result, we developed a novel setup based on a glass micro-prism to generate Evanescent Wave (EW) through Total Internal Reflection (TIR) microscopy. This prism-based design makes the system very compact, portable and low-cost. At the same time, a key advantage of measuring nanoparticle scattering is the high sensitivity of the analysis. The experimental setup was meticulously designed to house all optical elements within a custom-built metallic box, reducing its size to 40×30×40 cm^3^ and facilitating transport (figure 2a). The optical scheme of the prism-based TIR microscope, illustrated in Figure 2b, allows for significant size reduction compared to objective-based TIR microscopy setups, as reported in Table I. In particular, the illumination source consists of a 640 nm-diode laser with 10 mW power, collimated by a 20-mm lens (L1). Mirrors M1 and M2 are used to direct the excitation beam towards an ad-hoc glass micro-prism before it reaches the sample holder. M1 raises the excitation beam perpendicularly, while M2, fixed on the vertical wall of the metallic box, adjusts the inclination of the laser beam via a micrometre screw to achieve the incidence angle required for total internal reflection. The glass micro-prism is optically coupled to the glass coverslip through immersion oil, which has the same refractive index as glass (n_glass_ = n_oil_ = 1.516). Then, an iris positioned along the laser’s beam excitation path, in a plane conjugated to the image plane and the objective’s focal plane, spatially limits the excitation beam and regulates the illumination area. During measurements, the LOC-NPs sample is placed on a support atop the micro-prism, mounted on a 12.7 mm XYZ translation stage with standard micrometre screws, allowing spatial exploration of the sample and scanning measurements. After excitation, the scattered signal is collected by a CSI Plan Fluor 40X Nikon objective, which can be focused on the desired plane by moving the objective in the z-direction using a micrometre screw. The objective, characterised by a 0.6 numerical aperture (NA) and an extra-long working distance (2.8 - 3.6 mm), accommodates the distance constraints imposed by the glass micro-prism’s dimensions (Figure 2b, left inset). The collected signal is then transmitted to a CCD camera through mirror M3 and a tube lens (f = 200 mm). The tube lens and CCD camera are optically isolated from the environment by an 18 cm-long black tube housing an appropriate ND filter (with OD varying from 0.5 to 2.0) to attenuate the scattering signal and prevent camera saturation. Finally, the CCD camera is connected to a lab computer for image acquisition and analysis, explained in detail in section 2.3.

**Table I:** Comparison of LODs and technical characteristics between the traditional bulky setup and the compact TIR prototype.

| Feature | Traditional TIR Setup | Compact TIR Setup |
| --- | --- | --- |
| LOD | 1 ag/mL | 1 fg/mL |
| Dimensions | Standard optical table | 40x40x30 cm |
| Camera FOV | 42 $\mu\text{m}$ x 42 $\mu\text{m}$ | 221 $\mu\text{m}$ x 166 $\mu\text{m}$ |
| Camera type | EMCCD | CCD |
| Lateral resolution | 346 nm | 702 nm |
| Overall cost | $\sim 150\text{k €}$ | $\sim 10\text{k €}$ |

**Figure 2:**
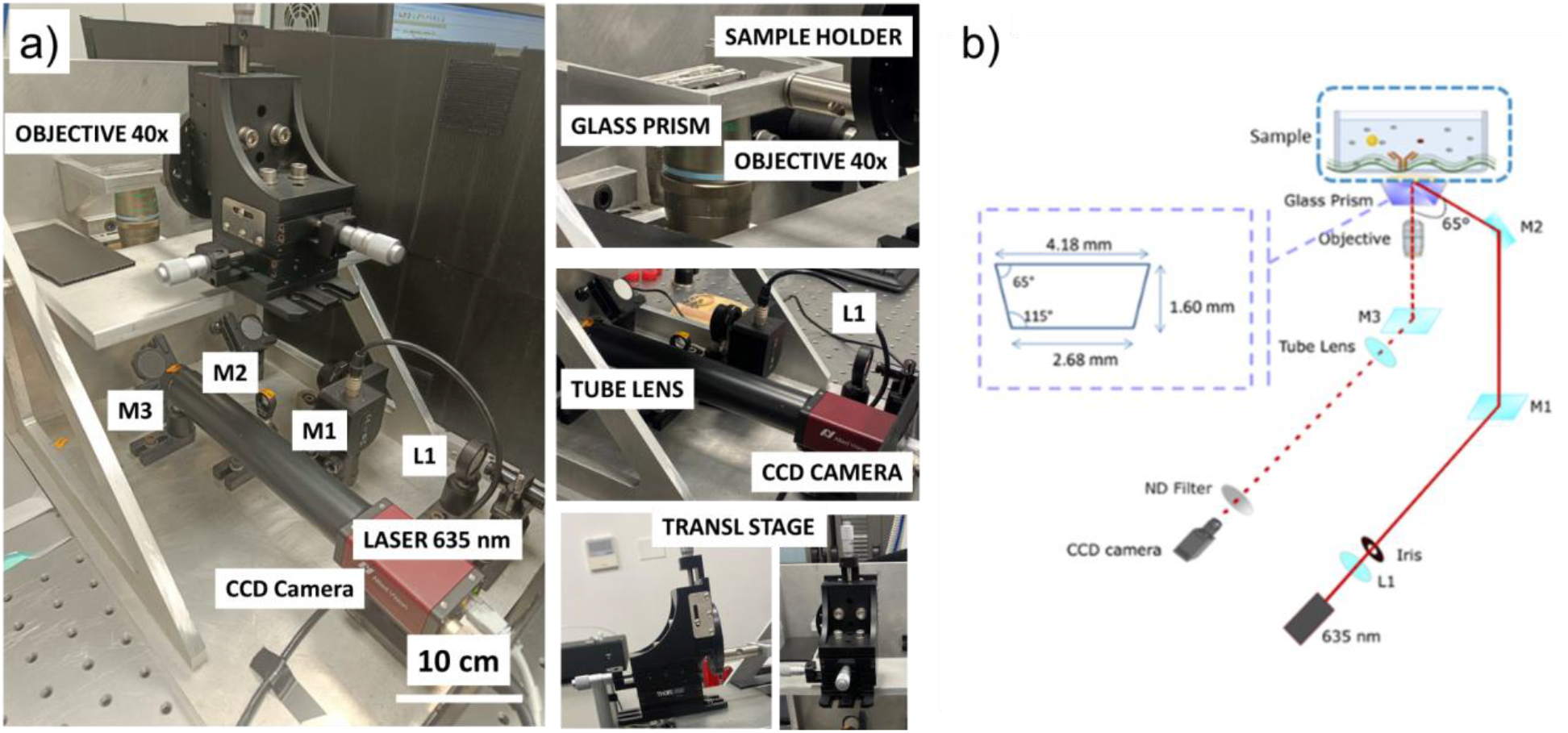
**(a)** Photos of the compact TIR setup from different views, showing the optomechanical components described in the main text. **(b)** Scheme of the optical path. L1: collimating lens (f=20mm); M1,2,3: Dielectric mirrors. Inset: geometry and dimensions of the glass microprism.

### 2.2. Synthesis of functionalized nanoparticles for the selective recognition of LPS

Functionalized nano-constructs were prepared starting with the synthesis of citrate-stabilised gold nanospheres (AuNSps) using the Turkevich-Frens synthetic protocol ^24^, further details are listed in section S1 in SI. An exchange ligand process was then implemented immediately after synthesis, following recent results obtained by our group ^21^, to functionalize the gold nano-surface with thiolated-peptide molecules designed to recognize bacterial lipopolysaccharides (LPS) changes in the serum of patients with severe infections or sepsis (Figure 3a) ^19^. The plasmonic spectra of the colloidal solution exhibited a characteristic resonance peak at approximately 520 nm for citrate-stabilised AuNSps, which shifted slightly to 522 nm upon peptide conjugation due to modification of the extinction coefficient (Figure 3b). Moreover, considering that the molecular weight of the peptide (5000 Da) is 25 times higher than citrate molecules (around 200 Da), peptide conjugation was also confirmed by changes in the size of AuNSps from 14±3 nm to 88±1 nm, as observed through autocorrelation curves and histograms in Figure 3c. Raman measurements further confirmed peptide conjugation to the AuNSps surface, with vibrational peaks indicative of amino acids becoming predominant due to the presence of the metallic nanostructures when peptide is attached to the NPs surface (Figure 3d).

**Figure 3:**
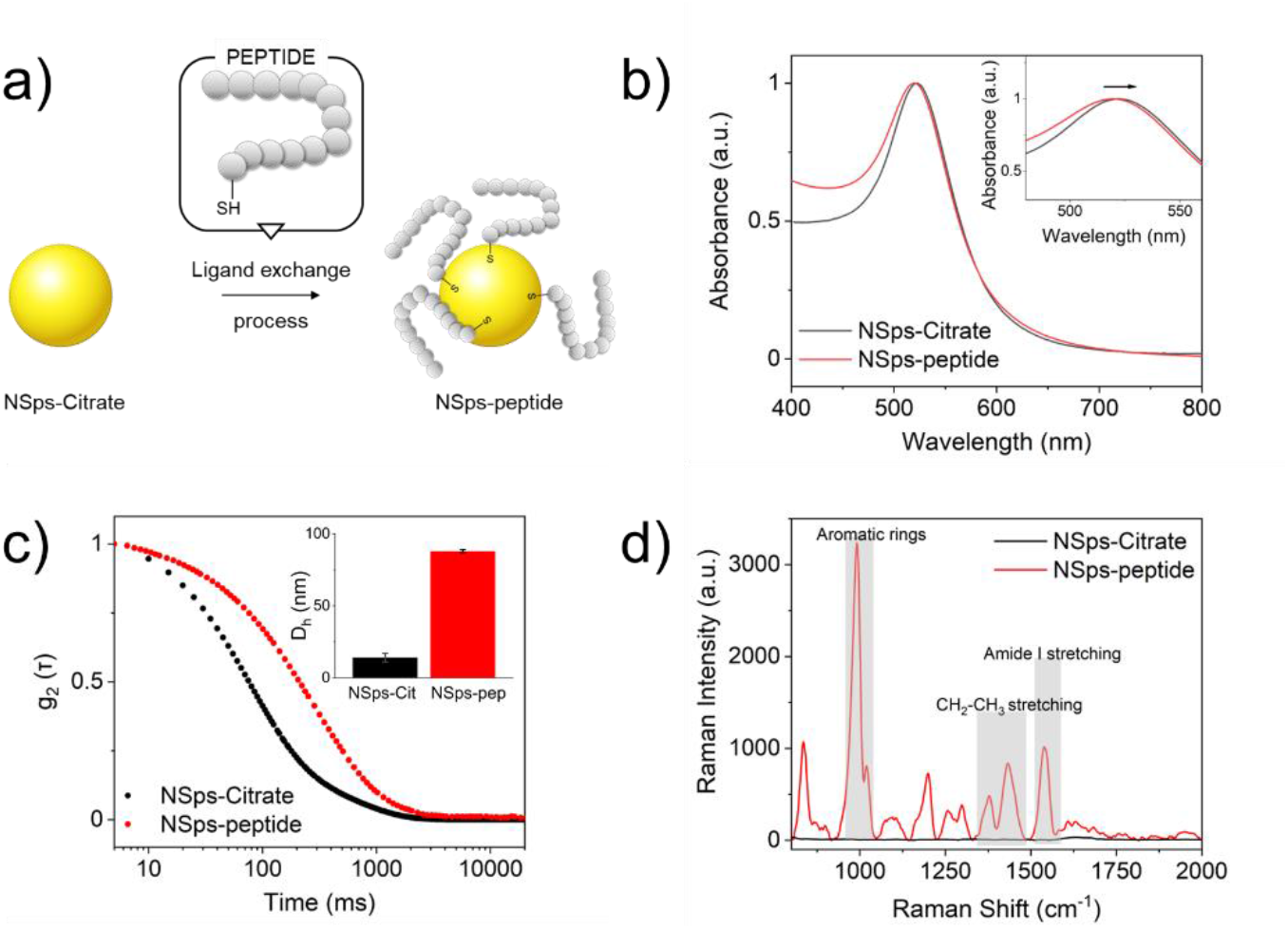
**(a)** Schematic procedure of gold nanoparticles functionalization with the thiol-functionalized peptide synthesis for the selective capturing of LPS. Characterization of NPs-peptide construct: **(b)** UV-Vis absorption spectra of NSps-citrate (black line) and NSps-peptide (red line). Inset shows the region 520-560 nm; **(c)** Autocorrelation curves of NSps-citrate (black line) and NSps-peptide (red line) measured through Dynamic Light Scattering analysis. Inset shows size distribution and ζ potential of NSps-citrate and NSps-peptide; **(d)** Raman spectra of NSps-citrate (black line) and NSps-peptide (red line). Measurements were performed with 785 nm laser, 20 mW, time acquisition 60s and 2 accumulations.

### 2.3. Image processing and analysis

The acquired images were analysed and post-processed through ImageJ by performing the procedure described in the scheme in Figure 4.

**Figure 4:**
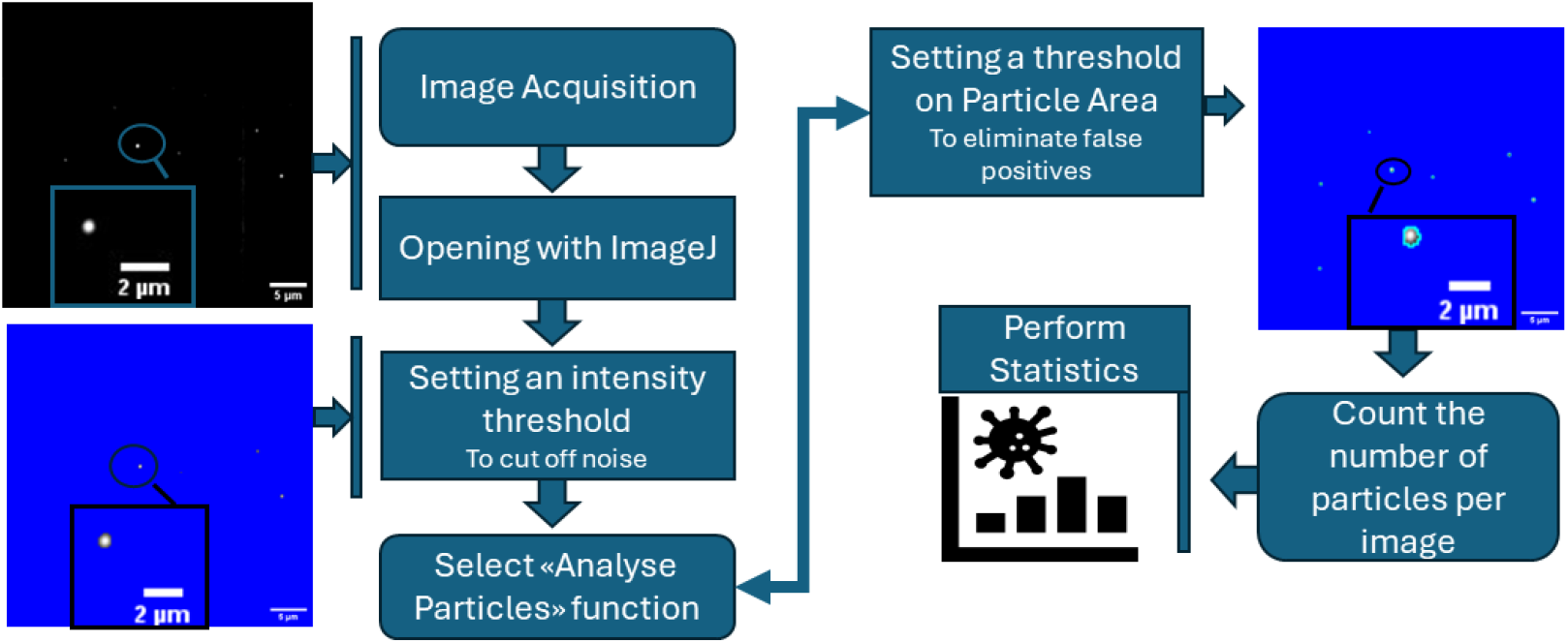
Graphical scheme of image processing procedures: first, the image is opened with ImageJ and a threshold value is chosen to remove the background noise. After noise subtraction, the function “Analyse particles” is exploited. By selecting this function, the user is asked to choose a minimum size value for particles, which was set to be of 10 pixel^2^. This value comes from the calculation of the lateral resolution of the image, which is 702 nm (see Supplementary Information section S7). After setting the thresholds on background and on particle area, the ImageJ function allows the user to select the quantities to be measured. In this way we obtain the number of particles detected, their surface areas and their average intensities for each image.

Samples preparation procedures and experimental parameters are reported in section S2 and S6 in SI. After image processing, statistical analysis on the obtained values was performed as follows:

i. 20 images were collected at each LPS concentration and were processed as depicted in Figure 4.
ii. The number of AuNPs detected in each image was divided by the image area (110 μm × 83 μm) to obtain the AuNPs density, d.
iii. The mean numerical density and its standard error were extracted from the densities values in (ii).
iv. For further statistical analysis, pair sample t-tests were conducted to determine if the mean AuNPs density at each concentration was statistically different from the mean density values obtained for the other concentrations of LPS..

## 3. RESULTS AND DISCUSSION

### 3.1. Validation of effective EW scattering of nanoparticles, LOD establishment and calibration curve determination

The capability of the compact EW-optical reader to detect LPS was tested by performing scattering measurements on gold nanoparticles functionalized for specific targeting. The results were then compared with the performances of the objective-based bulky setup, described in section “Validation of effective EW scattering of NPs and LOD establishment using the traditional bulky TIR Microscope” in section S4 of SI. Description of the traditional bulky setup is provided in section S3 in SI. LOC-NPs samples were dispersed in a PBS buffer at different concentrations, ranging from 1 ng/ml to 1 fg/ml, as described in section “LOC-NPs sample preparation” in section S2 in SI. The field of view (FOV) analysed corresponds to 110 μm × 83 μm, and N≥ 20 FOVs were collected for each LPS concentration. Image analysis and statistics on data were performed as described in section 2.3. Experimental results are reported in Figure 5a, showing that the density of NPs increases with increasing LPS concentrations, demonstrating the capability of the system to detect diverse LPS levels. Example images acquired at different LPS concentrations are shown in Figure 5b. These data were used to build the calibration curves, which equations are reported in section S8 in SI, correlating NPs density with LPS concentration (Figure 5c,d,e). The data points represent mean NPs density values with error bars indicating the standard error. Three different curves are shown: the one illustrated in Figure 5c shows the mean NPs density values obtained for all the values of LPS concentrations measured, ranging from 1 ng/mL to 1 fg/mL. The curve in Figure 5d focuses on the high concentrations range (10 pg/ml to 1 ng/ml), whereas the curve in Figure 5e zooms in on the low concentrations range (1 fg/ml to 10 pg/ml). A linear fit (red line) is applied to both these ranges, and the high R-square values indicate an excellent fit, suggesting a strong linear correlation between NPs density and LPS concentrations. Overall, the low-cost portable setup showed a very similar trend to the bulky setup when lowering LPS concentration (see section S4 in SI), with a limit of detection (LOD) of 1 fg/ml. The limit of detection was considered to be the lowest LPS concentration at which the mean NPs density was statistically distinguishable from the density of the control sample, in which LPS are not present. Although this LOD is three orders of magnitude higher than the bulk setup (1 ag/ml), likely owing to the less performing optical components chosen to reduce the cost and size of the system (Table I), the setup demonstrated improved sensitivity compared to other techniques currently in use in clinics, such as commercial ELISA kits, which report limits of detection in the tens or hundreds of picograms per mL range ^25,26^.

**Figure 5:**
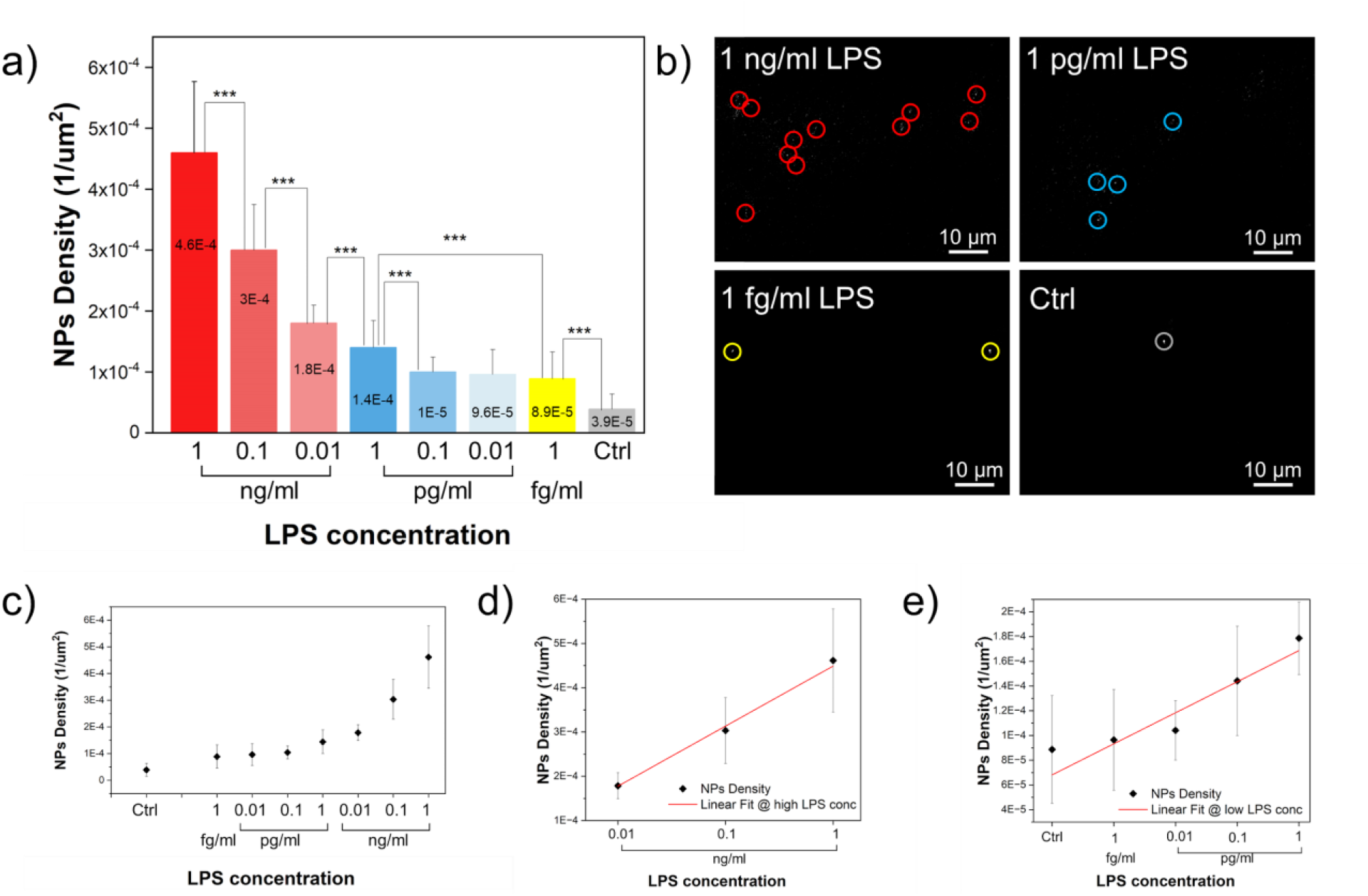
**(a)** Graph showing the results obtained by using the compact TIR prototype. Scattering measurements were performed at LPS concentrations spanning from 1 ng/mL down to 1 fg/mL. N≥ 20 for each LPS concentration. NPs density values are illustrated together with the associated standard errors (error bars). Pair student t-tests have been performed to evaluate the statistical differences between different LPS concentrations. The LOD is set at 1 fg/mL. **(b)** Scattering images at different LPS concentrations acquired with the compact optical setup. Bright circled spots are scattered AuNPs. Scalebar 10 μm. **(c)** Linear-logarithmic/semilog Graph of NPs mean densities in function of LPS concentrations. **(d)** Linear-logarithmic semilog graph of NPs mean density in function of low concentrations of LPS (< 10 pg/mL). Red line represents the fitting equation. **(e)** Linear-logarithmic graph of NPs mean density in function of high concentrations of LPS (≥ 10 pg/mL). Red line represents the fitting equations, described in section S8.

### 3.2. Reproducibility and Robustness Assessment

Once the capability of the compact prototype to detect highly diluted analytes in aqueous solutions has been validated and the LOD of the system established, tests were performed to characterise the reproducibility of EW scattering experiments and to test the reliability of the entire system. To this end, four different LPS concentrations in the range of 1 ng/mL - 1 fg/mL were chosen, five samples were prepared for each concentration and 24 images were acquired for each sample. NPs density was calculated and a mean NPs density was extracted along with the corresponding standard error. Afterward, the average on mean densities was calculated over the five samples of each concentration to extract a final mean density and its standard deviation as reference value (red bars in Figure 6). Further statistics was done by performing pair sample t-student tests to establish whether the mean NPs densities of different LPS concentrations showed significant statistical differences. Results indicate that densities are different from each other at the 0.05 level (p<0.05). The histograms in Figure 6 show the results, where the white bars represent the NPs density extracted from the five samples at each LPS concentration, along with the associated error bars representing the standard error of the mean of the distribution. Reproducibility tests showed consistent results across different LPS concentrations, proving the reliability of the compact prototype in performing accurate measurements.

**Figure 6:**
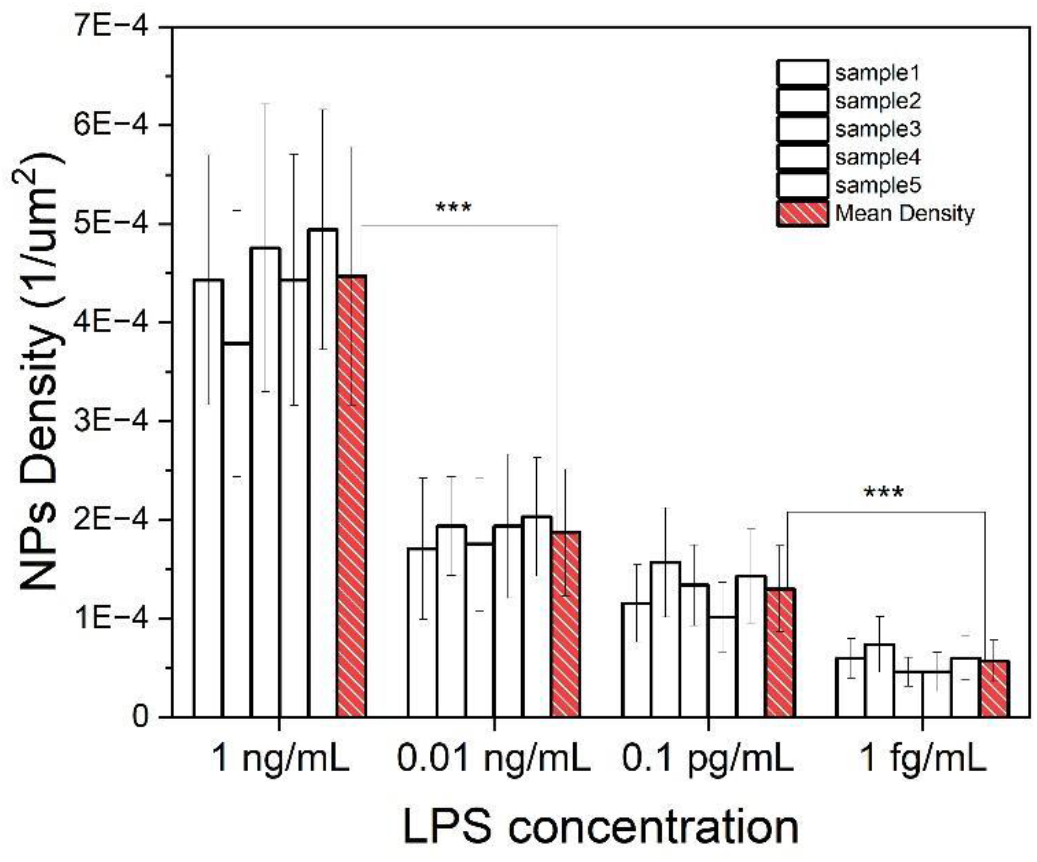
Histogram showing the reproducibility measurements performed on four different LPS concentrations. 5 different LOC-NPs samples were observed for each concentration (white bars), (N=24 for each concentration) and the mean value of their density distribution was calculated, along with its standard error. Red bars represent the mean density values calculated from the 5 samples with the same LPS concentration, error bars are standard errors.

### 3.3. Validation on human samples

Finally, the capability of the compact setup was further tested on human serum samples enriched with known concentrations of LPS. Serum vials derived from healthy patients were provided by the Department of Medical Biotechnology of the University of Siena, and were used to perform measurements. Serum samples spiked with three different concentrations of LPS (1 ng/mL, 1 pg/mL, and 1 fg/mL) were measured. The setup successfully detected LPS concentrations down to 1 fg/ml. Calibration curves calculated from the fitting (Figure 5d,e) were used to calculate the expected NPs density values through equations (1) and (2) (SI section S8), and these were then compared with the actual values obtained from measurements in spiked serum, to quantify the recovery rates (RR) at each LPS concentration. RR was obtained by dividing the mean NPs density value measured in spiked serum samples by the expected density value calculated through the fitting equations (1) and (2) of the calibrations curves. All these values are listed in table in figure 7b. Figure 7a illustrates the expected values obtained through fitting equations (black bars) compared with real densities obtained from measurements (red bars), as well as the points obtained in aqueous solution (blue bars) at the same LPS concentrations. Comparison with measurements in aqueous samples showed no significant difference, highlighting the system’s capabilities to handle human biofluids effectively.

**Figure 7:**
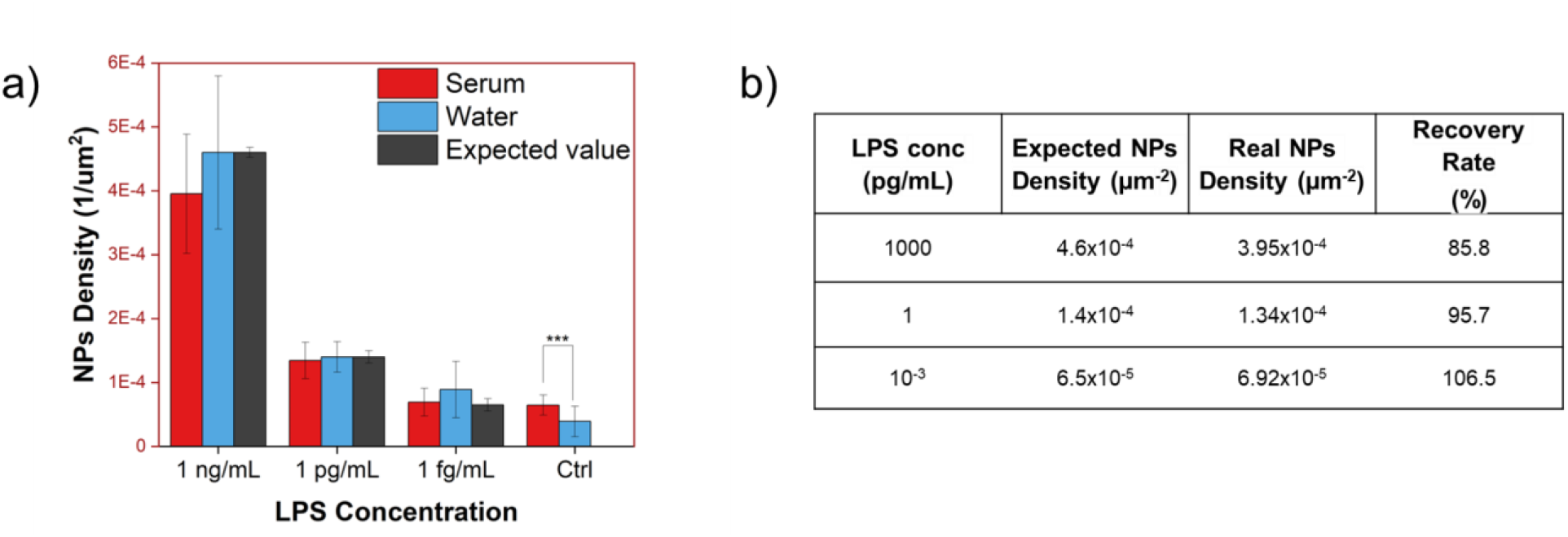
**(a)** Histogram showing the mean NPs density values obtained from experiments in human serum (red bars, N>60), in synthetic aqueous samples (blue bars, N>20) and calculated through the fitting equations (black bars) at specified LPS concentrations. Student t-tests have been done to determine if there were significant differences between different samples at the same concentration of LPS, the only difference being in the control sample, as expected. **(b)** Table comparing the measured and expected mean density values at specified LPS concentrations and the corresponding Recovery Rate (RR).

Notably, a significant difference between density values in water and in serum was observed in the control sample (Figure 7a). This result was expected since it is known that a small amount of LPS is present in the serum of healthy subjects. Therefore, when no LPS were added to the serum solution, a higher NPs density was observed compared to the aqueous control sample, where LPS were not present. This finding is important because it underscores the ability of our custom-made setup to detect the basal concentration of LPS in healthy patients, even though the corresponding NPs density value is not statistically distinguishable from the one of the 1 fg/mL serum spiked sample.

## 4. CONCLUSIONS

In conclusion, our study highlights the advantages of combining confined evanescent wave (EW) excitation, with highly efficient scatterers such as gold nanoparticles, as a robust methodology for high sensitive detection of target analytes in liquid samples. The integration of this approach with a compact and portable total internal reflection (TIR) microscope further demonstrates its potential for the clinical scenario. Our investigation thoroughly evaluated the limit of detection and reproducibility of this compact setup, characterised by reduced dimensions and constructed using cost-effective optomechanical components. Notably, this user-friendly and easily reproducible system exhibits the capability for rapid, accurate, and multiplexed analysis. As a proof of concept, we applied the device to the detection of bacterial lipopolysaccharides (LPS) in human serum, a key indicator of potential sepsis. The achieved limit of detection, set at 1 fg/mL LPS concentration, stands as a remarkable accomplishment, especially when considering the limited cost of the system and its competitive performance against established techniques. Given the versatility of nanoparticles (NPs) engineering, our proposed device emerges as a versatile tool for the real-time analysis of various diseases in human biofluids. Its compact design, ease of use, and affordability makes it a promising candidate for widespread application in clinical settings. The successful validation using human serum from patients, coupled with its ability to detect ultra-low concentrations of target analytes, pave the way for its potential application in mass population screening, early-stage disease diagnosis and bed-monitoring.

## AUTHOR INFORMATION

### Author Contributions

The manuscript was written through contributions of all authors. All authors have given approval to the final version of the manuscript. ‡These authors contributed equally.

### Notes

The authors declare no competing financial interest.

## ACKNOWLEDGMENT

This work was supported by FAS Regione Toscana under the DIVISA project – Progetti di alta formazione attraverso l’attivazione di AR - and by the “Integrated infrastructure initiative in photonic and quantum sciences – I-PHOQS” project finances by the EU next generation PNRR action. The authors would like also to thank the “Centro di competenza “RISE” funded by FAS Regione Toscana.

---

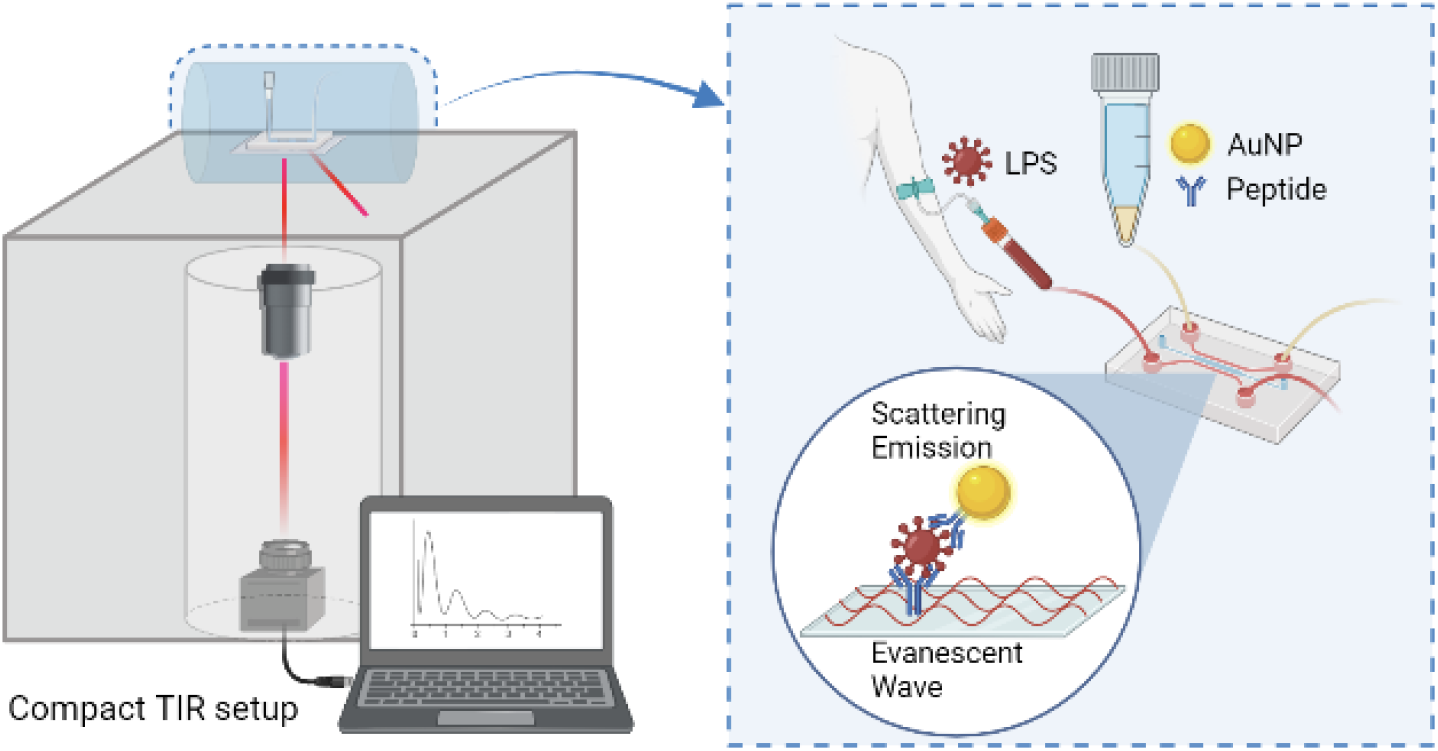

