## Supplementary Information for "A compact prism-based microscope for high sensitive measurements in fluid biopsy"

*S1	Nanoparticles synthesis and functionalization*

Gold-nanoparticles (AuNPs) were synthesized exploiting the seeded-growth process described by Yuan et al [14]. The seed solution of 15 nm-nanospheres (AuNSps) was prepared by adding 1.5 mL of 30 mM HAuCl4·3H2O (1%) to 48.5 mL of boiling and stirring MilliQ (750 rpm, 250°C). After 10 s, 4.5 mL of 38.8 mM sodium citrate solution were added to the solution. The boiling solution was stirred under heating for 15 min, and then cooled at room temperature under magnetic stirring for 30 min. Later, a nonnatural antimicrobial peptide was used in a ligand‐exchange reaction to replace the citrate layer, in order to exploit its specific capacity to target lipopolysaccharide (LPS) from gram-negative bacteria ^1^. The conjugation exploits the thiol pendant moieties characteristic for the synthetic peptide. In particular, 2.5 μL of a 2.5 μM aqueous solution of peptide were added to 500 μl of 5 nM solution of AuNSps and stirred at room temperature for at least 2 hours to reach [peptide]/[AuNP] molar ratio of 2.5. The concentration of AuNSps was determined according to the formula reported in the method of Liu et al. ^2^. Peptide-capped nanoparticles (NSps-peptide) were then centrifuged in 1.5 mL Eppendorf tubes at 25 °C, 10 000 rpm for 40 minutes and redispersed in milliQ water. The plasmonic properties of gold NPs colloidal solution before and after functionalization processes were acquired in the range from 400 nm to 900 nm with a UV-Vis‐NIR spectrophotometer (Lambda 950 instrument, Perkin Elmer, Waltham, MA, USA). This instrument uses a deuterium and tungsten halogen lamp as the light excitation source. UV WinLab Software (Perkin Elmer, Waltham, MA, USA) is used to acquire spectra and data is processed with Origin software. The hydrodynamic dimensions were characterised by Dynamic Light Scattering (DLS) analysis performed with a Malvern Zetasizer Nano series ZS90 (Malvern, Worcestershire, UK). Measurements were performed with a fixed scattering angle of 90°, at 25°C. Each sample was measured three times and each measurement consisted of about 30 acquisitions. Cumulating statistics were used to measure both the hydrodynamic diameter and the polydispersity.

*S2	LOC-NPs Samples preparation*

After NPs functionalization, it was necessary to verify that effective EW scattering occurred when using the LPS-peptide-NPs sandwich, and this was tested on the objective-based bulky TIR setup. Custom-made fluidics channels were created by attaching glass coverslips on top of microscope slides by means of three parallel strings of  double-sided tape. This procedure allows us to obtain two different channels for each microscope slide used. Liquid samples can be easily fluxed into these chambers by dropping a few microliters of the solution at one side of the channel, and a second fluid can later be fluxed by exploiting the capillarity effects of the channels. LOC-NPs samples to be coupled with the traditional bulky TIR microscope for the analysis of scattering signals coming from gold nanoparticles were prepared as follows: 30 𝜇L peptide 1 mg/mL were fluxed within the chamber and incubated for 5 minutes; this is followed by 3x washing with PBS 1X. Then, 30 𝜇L LPS solution at the desired concentration (ranging from 1 ng/ml to 1 ag/ml) were fluxed within the chamber and incubated for 4 minutes, followed again by 3x wash with PBS 1X. 30 𝜇L NPs-pep 0.14 nM were fluxed within the chamber and incubated for 4 minutes, then the samples were 3x washed with PBS 1X (Figure S1). Samples were prepared starting from a LPS concentration of 1 ng/mL and then decreasing the concentration of one order of magnitude at each new sample, until the final concentration of 1 ag/mL. A reference control sample was prepared by following all the steps described above but avoiding the step of fluxing the LPS solution in the chamber. To perform scattering measurements, LoC-NPs devices were placed on the sample holder of the objective-based TIR microscope.

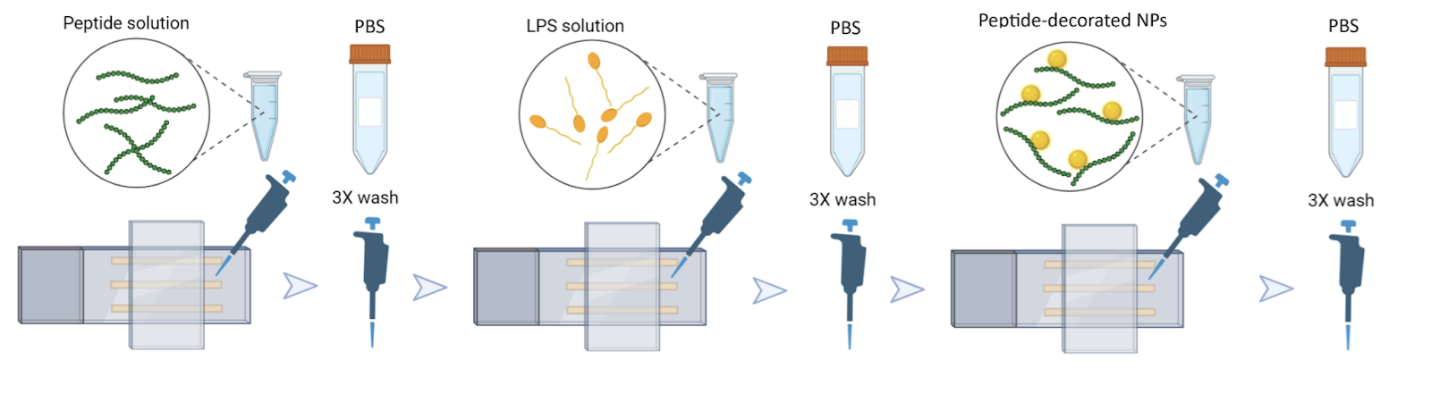

**Figure S1:** Schematic representation of the preparation steps of samples. After creating two channels on the microscope glass slide, the solutions are fluxed into the channels in the order described in the figure, the incubation time being 5 minutes at the first step, and then 4 minutes for the following steps.

*S3	Description of the traditional TIR setup*

The optical setup is composed of an inverted wide-field fluorescence microscope (Nikon ECLIPSE TE300) equipped with a 643 nm diode laser source for fluorescence and scattering excitation ^3–5^. The laser beam is then magnified 10 times by an achromatic doublet telescope L1,L2. The excitation beam is reflected by the mirror M, after a filtration through a variable beam stop, and focused onto the back focal plane of the objective (Nikon 60X, oil immersion, NA 1.49 TIRF) by lens L3. The mirror M and the lens L3 are both mounted on a motorized translator to displace the excitation beam laterally at different distances from the optical axis of the objective and change between epifluorescence, highly inclined, and TIR optical configurations. The excitation laser beam is directed along the optical axis of the objective through a 50:50 Beam Splitter (50:50 BS). The scattered light is then collected through the same 50:50 BS and physically filtered through an iris housed in the dichroic filter cube below the objective to stop reflections at the optics edges that could strongly affect the detection of the light scattered by single emitters. The collected signal is then projected onto the EMCCD camera (Andor iXon X3) by the tube lens L4, followed by a 3X telescope (L5, L6). Before reaching the camera, the scattered collected light is filtered by an ND filter. Full field of view is 42 × 42 µm2 wide, with 91 nm pixel size.

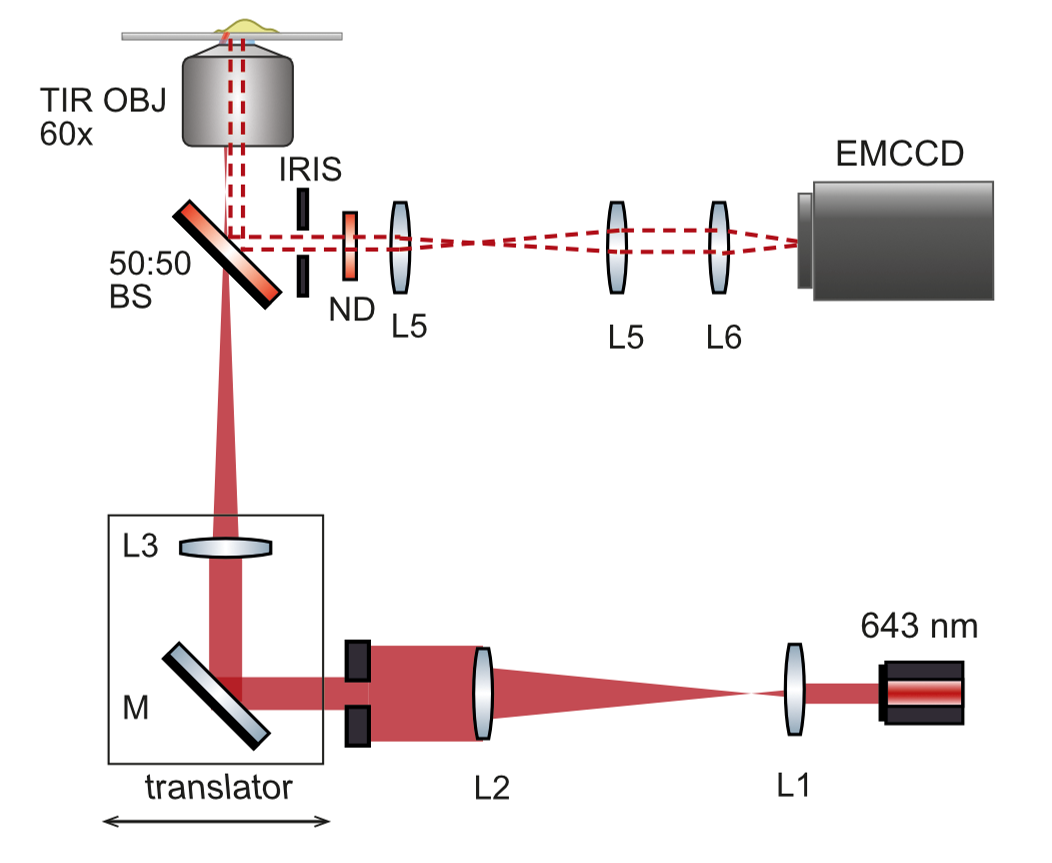

**Figure S2.** Optical scheme of the bulk TIR setup.

*S4	Validation of effective EW scattering of NPs and LOD establishment using the traditional bulky TIR microscope*

To validate the efficiency of the NPs-based biosensor in detecting LPS with high sensitivity, we performed scattering measurements on a state-of-the-art objective-based TIR microscope developed in our laboratory, with single molecule sensitivity and nanometer localization precision. The imaging setup is shown in Figure S2. LOC-NPs samples were prepared as described in section S2, starting with an LPS concentration of 1 ng/mL and decreasing it by one order of magnitude for each new sample. Images were collected and analyzed as described in section 2.3. All the images reported here have been acquired with an exposure time of 500 ms. The laser beam is characterized by an excitation spectrum peaked at 640 nm, with an output free-space power of 13 mW. The image field of view is (42 μm × 42μm). N= 20 FOVs were randomly acquired for each sample. Results are shown in Figure S3, along with representative scattering images at different LPS concentrations. As expected, the calculated density of NPs detected through EW scattering decreased as the LPS concentration diminished. Notably, we detected NPs down to 1 ag/mL, setting this as the LOD of the bulky system, since it was the lowest LPS concentration producing a statistically different density from the control sample.

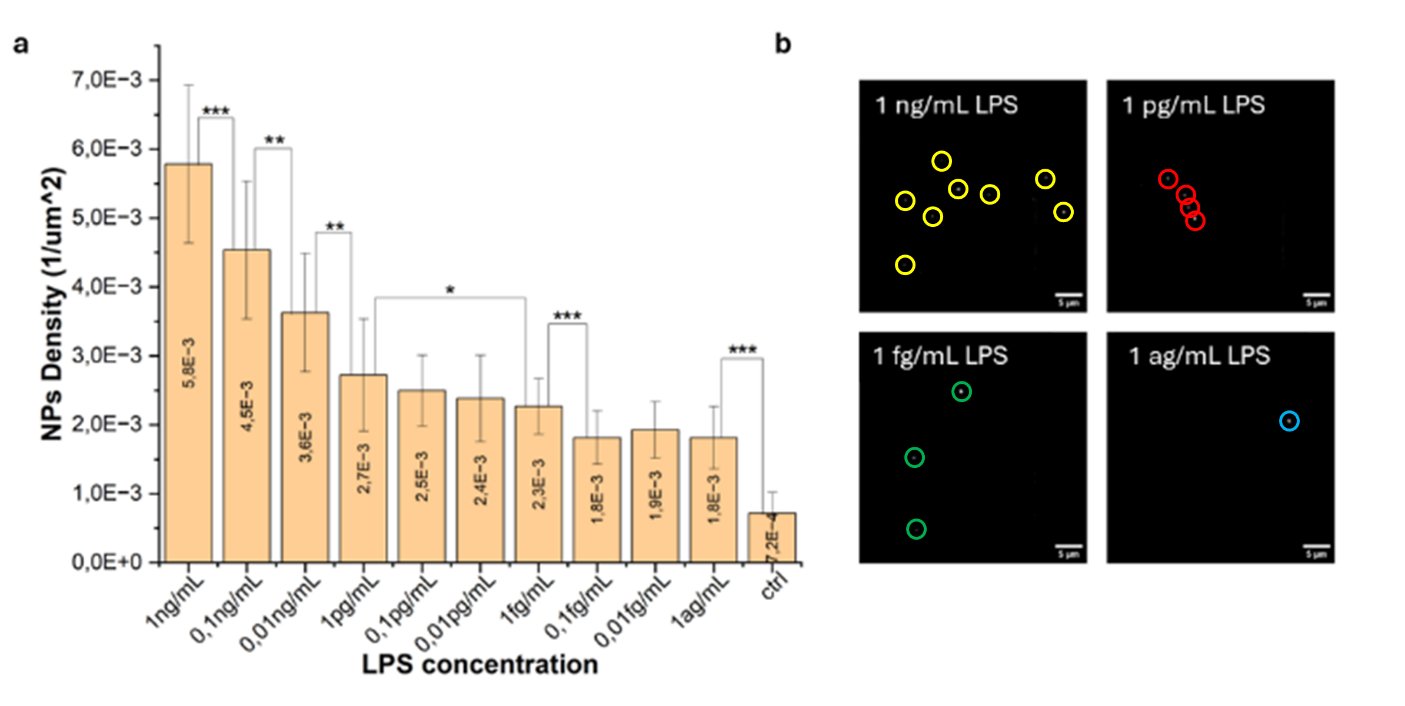

**Figure S3.** (a) Graph showing the results obtained by using the bulky TIR microscope. Scattering measurements were performed at LPS concentrations spanning from 1 ng/mL down to 1 ag/mL. N= 20 for each LPS concentration. NPs density values are illustrated together with the associated standard errors (error bars). Pair student t-tests have been performed to evaluate the statistical differences between different LPS concentrations. The LOD is set at 1 ag/mL. (b) Scattering images at different LPS concentrations acquired with the bulky microscope. Bright circled spots are scattered AuNPs. Scalebar 5 μm.

### *S5 Lateral image resolution in the traditional TIR setup*

The value of 5 pixel2 which was used as threshold value for particle dimensions in image analysis comes from the calculation of the lateral resolution of the image. The image lateral resolution was calculated by choosing a NPs which was clearly distinguishable and not agglomerated with other particles and by drawing an horizontal line crossing the particle (Figure S4a). With the ImageJ function called “Plot profile”, we can plot the intensity gray values in function of distance from the starting point of the drawn line. By performing a Gaussian Fit on the data resulting from this plot, we can obtain the FWHM of the gaussian function, which corresponds to the PSF diameter. In this case, it resulted to be of 346 nm (Figure S4b). The mean PSF radius is then $\frac{346}{2} nm$and the mean particle area, if we consider a squared region, is $\left( \frac{346}{2} \right)^{2}nm^{2}=29929 nm^{2}.$ Since the pixel area is $\left( 82 nm \right)^{2}=6724 nm^{2},$when setting a threshold on the particles dimensions, the pixel area must be multiplied for a factor given by

$$\frac{Mean particle area}{Pixel area}=\frac{29929 nm^{2}}{6724 nm^{2}}= 4,45$$

Since this value takes into account the mean particle area, the value 5 was taken as an excess value.

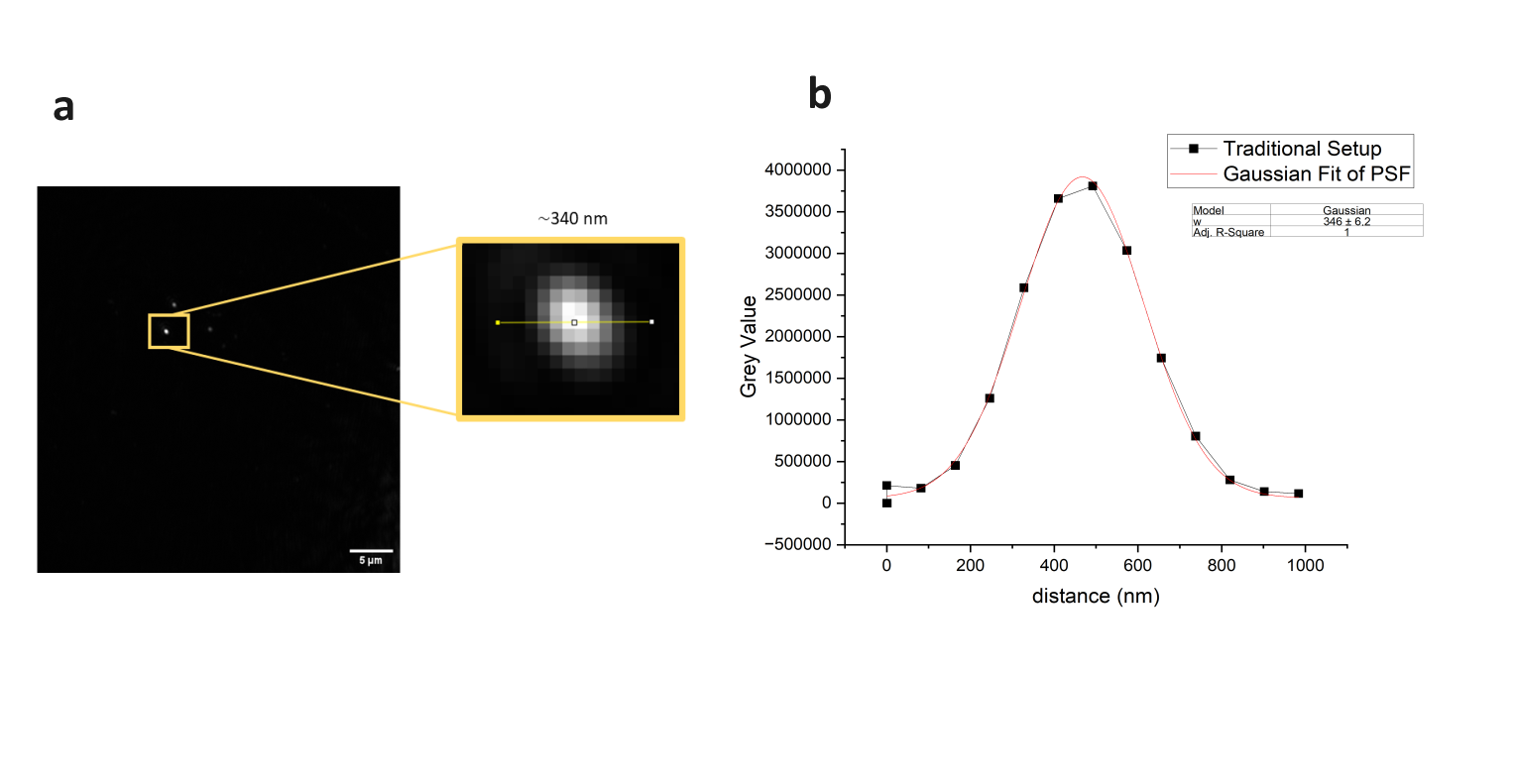

**Figure S4**: (a) Image acquired with the traditional TIR setup and zoom of the nanoparticle used to plot the PSF profile. (b) PSF profile of the gold nanoparticle and its gaussian fit. The FWHM (w value) of the Gaussian function provides the image lateral resolution.

### *S6 LOC-NPs Sample preparation procedures and Experimental Parameters for measurements performed with the compact TIR setup*

The novel bench-top prototype was tested against synthetic models and real serum samples. The former were prepared with the same procedure used for the ones measured on the bulky microscope. LOC-NPs samples to be coupled with the compact low-cost TIR prototype for the analysis of scattering signals coming from gold nanoparticles were prepared by fluxing solutions as described above. The highest LPS concentration was set again at 1 ng/mL, and it was decreased each time by one order of magnitude as before, A reference control sample was prepared as described before. The samples with real human serum were prepared by using serum vials derived from healthy patients that were provided by the Department of Medical Biotechnology of the University of Siena, by following these steps: serum collected from healthy people was spiked with the addition of 1 mg/mL LPS solution at various volumes to reach the final LPS concentration of interest (1 ng/mL, 1 pg/mL, 1 fg/mL).Once the samples are ready, the optical coupling between the glass-prism and the coverslip is established by spilling a drop of oil (n=516) on top of the glass microprism. The LOC-NPs sample is then positioned on the custom-built holder. Then, the incidence angle of the excitation beam is adjusted to obtain the EW, which will propagate through the aqueous solution of the sample within the characteristic distance of 200 nm.

All the measurements reported here have been performed with a 1.5 ND filter in the filtering unit before the CCD camera. All the images have been acquired with an exposure time of 500 ms. The laser beam is characterised by an excitation spectrum peaked at 638 nm, and its power was measured to be of 9.9 mW before passing through any optical element. The image field of view is (110μm × 83μm) . N≥20 FOVs were acquired for each sample for the determination of the LOD. For experiments on reproducibility, 4 LPS concentrations were tested, and 5 samples were prepared for each LPS concentration. For each sample, N=24 FOVs were acquired. For scattering measurements on human serum samples, LPS concentrations of 1 ng/mL, 1 pg/mL and 1 fg/mL were observed, as well as the reference control sample, and N≥60 FOVs were acquired for each concentration.

### *S7 Lateral image resolution in the compact TIR setup*

The value of 10 pixel2 which was used as threshold value for particle dimensions in image analysis comes from the calculation of the lateral resolution of the image. The image lateral resolution was calculated as explained in the previous section by choosing a NPs which was clearly distinguishable and not agglomerated with other particles and by drawing an horizontal line crossing the particle (Figure S5a). With the ImageJ function called “Plot profile”, we can plot the intensity gray values in function of distance from the starting point of the drawn line. By performing a Gaussian Fit on the data resulting from this plot, we can obtain the FWHM of the gaussian function, which corresponds to the PSF diameter. In this case, it resulted to be of 702 nm (Figure S5b). The mean PSF radius is then $\frac{702}{2} nm$and the mean particle area, if we consider a squared region, is $\left( \frac{702}{2} \right)^{2}nm^{2}=123201 nm^{2}.$ Since the pixel area is $\left( 114 nm \right)^{2}=12996 nm^{2},$when setting a threshold on the particles dimensions, the pixel area must be multiplied for a factor given by

$$\frac{Mean particle area}{Pixel area}=\frac{123201 nm^{2}}{12996 nm^{2}}= 9,5$$

Since this value takes into account the mean particle area, the value 10 was taken as an excess value.

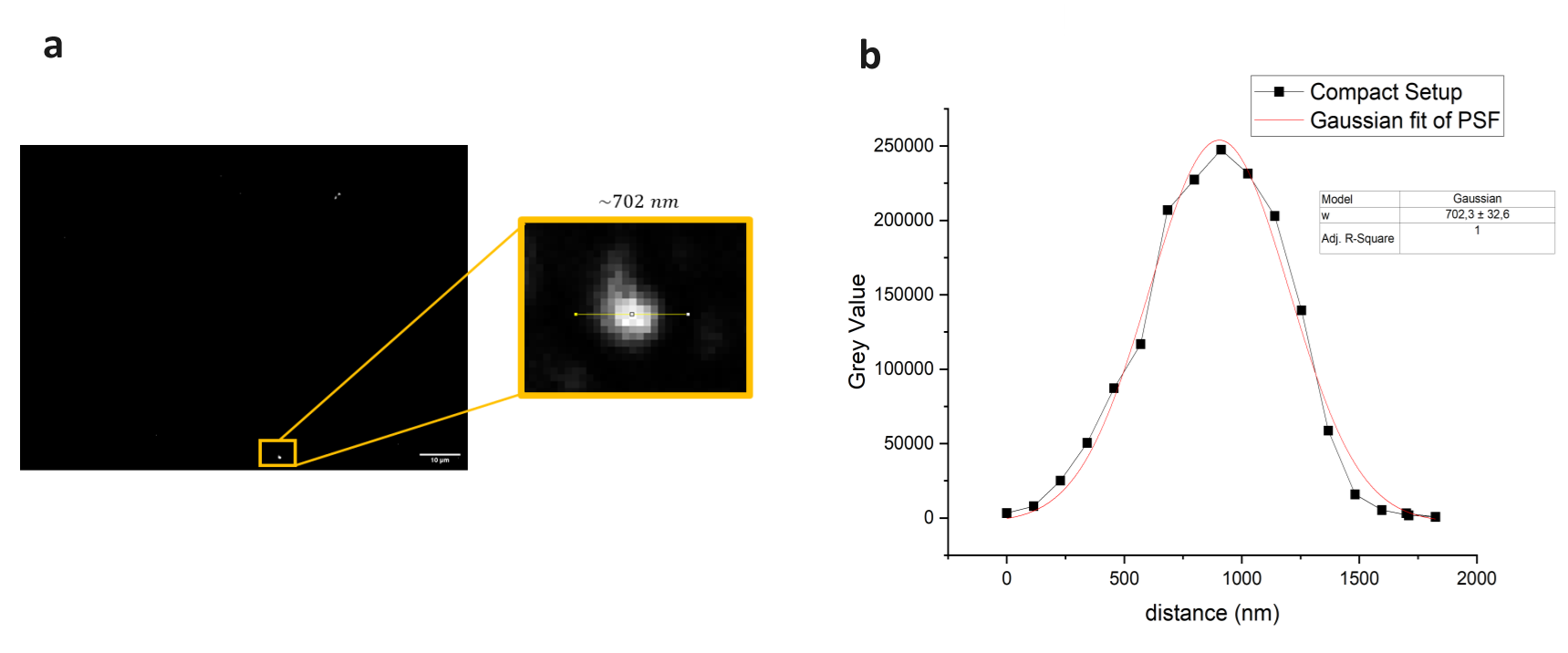

**Figure S5**: (a) Image acquired with the compact TIR prototype and zoom of the nanoparticle used to plot the PSF profile. (b) PSF profile of the gold nanoparticle and its gaussian fit. The FWHM (w value) of the Gaussian function provides the image lateral resolution.

### *S8 Calibration curves for Recovery Rate calculation*

The functions describing the calibrations curves are:

Eq. (1):	$y [{\mu m}^{-2}]=4,2\times{10}^{-5}+(1,4\times{10}^{-4}){Log}_{10}(x [pg/mL]) \pm(9,7\times{10}^{-6})$;

Eq. (2):	$y [{\mu m}^{-2}]=1,4\times{10}^{-4}+(2,5\times{10}^{-5}){Log}_{10}(x [pg/mL])\pm(7,8\times{10}^{-6})$.

where $y$ is the NPs density expected value and $x$ is the concentration of LPS expressed in pg/mL.
